# The effect of altitude on the expression of immune-related genes in Peruvian rural indigenous

**DOI:** 10.1101/2024.03.06.583674

**Authors:** Luis Jaramillo-Valverde, Gilderlanio Santana de Araújo, Julio A. Poterico, Catalina Martinez-Jaramillo, Vicky Roa-Linares, Sandra Alvites-Arrieta, Nelis Pablo-Ramirez, Milward Ubillus, Diana Palma-Lozano, Rafael Tou, Carolina Silva-Carvalho, Luca Vasconcelos-da-Gama, Lucas F Costa, Manuel Corpas, Eduardo Tarazona-Santos, Soumya Raychaudhuri, Heinner Guio

## Abstract

**Background:** Genetic factors influencing immune response pathways in Andean populations may underlie adaptations to high-altitude environments. To investigate transcriptomic signatures associated with altitude, we analyzed immune-related gene expression across individuals residing at different elevations.

**Methods:** We recruited 62 Peruvian volunteers, predominantly from rural regions with high proportions of indigenous ancestry, living at low and high altitudes. Peripheral blood mononuclear cells (PBMCs) were stimulated with bacterial lipopolysaccharide (LPS), Pam3CSK4 (a synthetic triacylated lipopeptide), and R848 (an imidazoquinoline analog of viral nucleic acids). Population structure and ancestry were characterized, and transcriptome-wide differential expression analyses were performed.

**Results:** We identified 30 genes with significant altitude-associated expression differences, including 22 downregulated (e.g., *FN1, CD36, FOS*) and nine upregulated genes. Functional enrichment indicated roles in acute inflammatory response, leukocyte migration, and positive regulation of myeloid leukocyte differentiation.

**Conclusions:** High- and low-altitude Andean individuals exhibit distinct immune gene expression profiles, defining a population-specific transcriptomic signature that may reflect altitude-related immune adaptation.

**AUTHOR SUMMARY:** Andean populations exhibit genetic adaptations in immune response pathways, potentially linked to high-altitude living. To investigate this, we conducted genome-wide and transcriptome-wide analyses to identify immune-related gene expression differences between high- and low-altitude residents. We analyzed the genetic structure and ancestry of Peruvian individuals (primarily rural, with strong indigenous ancestry) living at different altitudes. Peripheral blood mononuclear cells (PBMCs) from 62 volunteers were stimulated with bacterial (LPS, Pam3CSK4) and viral (R848) mimics to assess immune responses. Differential expression analysis identified 22 down-regulated genes (e.g., *FN1, CD36, FOS*) and nine up-regulated genes, enriched in acute inflammatory response, leukocyte migration, and myeloid leukocyte differentiation. Our findings reveal a unique immune gene expression profile in Andean highlanders, distinct from other populations, suggesting adaptive modulation of both innate and adaptive immunity in response to high-altitude challenges.

## INTRODUCTION

Approximately 1% of the world’s population resides in high-altitude areas, specifically at elevations over 2500 meters above sea level (m.a.s.l.). Countries such as Peru and Bolivia have a significant portion (30 percent) of their population living at high altitudes in the Andean mountains (1). Living in high-altitude regions poses several environmental challenges for humans, including reduced oxygen levels, increased exposure to ultraviolet radiation, cold temperatures, and altered responses to pathogens, among other factors (2). Pioneering researchers in the field of population genomics at high altitude have suggested that genes of the Hypoxia Inducible Factor (HIF) pathway have evolved under positive natural selection (3,4). In contrast, the genes *ENDRA, PRKAA1*, and *NOS2A* have been suggested as being associated with blood pressure regulation and reproductive success in Andeans (5). Additionally, genetic studies have demonstrated that native Andean highlanders depict positive selection of cardiovascular genes, such as *BRINP3, NOS2, TBX5 (6)*, and the non-coding region *HAND2-AS1* (7).

As previously mentioned, hypoxia impairs oxidative metabolism, which forces cells to rely on oxygen-independent processes like glycolysis to produce ATP. Hypoxia plays a crucial role in immune cell metabolism regulation, which serves as a major regulator of immunometabolism, in addition to controlling immune cell phenotypic and function (8,9). Numerous studies have shown that hypoxia regulates cell types involved in the innate and adaptive immune response, including epithelial cells, neutrophils, macrophages, dendritic cells, T cells, B cells, natural killer cells, and innate lymphoid cells, since the first report of a role for HIF in the control of macrophage function in 2003 (10). A recent study confirms these findings and provides further insights. Analyses of a genome-wide study involving 286 individuals across Peru and over-representation functional analysis identified ten positive natural selection signals among highlander populations related to cardiovascular function (*TLL1, DUSP27, TBX5, PLXNA4, SGCD*), the hypoxia-inducible factor pathway (*TGFA, APIP*), skin pigmentation (*MITF*), glucose (*GLIS3*), and glycogen metabolism (*PPP1R3C, GANC*) (11), as well as genes involved in the *TP53* pathway, such as *SP100, DUOX2*, and *CLC (12)*.

These studies indicate that Peruvian Andean populations have exhibited distinct adaptations compared to other groups like Tibetans, Sherpas, or Ethiopians in response to high-altitude living (11,12). Moreover, specific genes related to immune response have been identified in Andean populations as being positively selected. These genes might provide insights into the adaptation of the immune system and responses to pathogens among those residing at high altitudes. Examples of robust adaptation signals related to altitude in Peruvian and Bolivian populations can be found in the studies conducted by Borda et al. (7) and Jacovas et al. (12). Specifically, these studies identified a significant association with the *DUOX2* gene, which plays a crucial role in maintaining homeostasis in thyroid hormone production, as well as influencing the innate immune system and inflammatory response.

Likewise, Caro-Consuegra et al. found positive selection in Peruvian Andean populations in *CD40, TNIP1*, and *CHIA* genes, which are involved in the innate immune system (11). Furthermore, genes such as *ANXA1, CD44, ADGRE1, KCNA3*, and *FCRL4*, among others, seem to have been positively selected due to *Mycobacterium tuberculosis* in Andean Ecuadorian populations, and this event could have emerged before the European contact with aboriginal communities (13). However, the aforementioned genomic studies have not included functional analysis to correlate genomic findings with other biomarkers and phenotypes.

To our knowledge, there is currently no investigation assessing the expression of immune genes in native participants (*e*.*g*., Andean Peruvians) with long-term exposure to high altitude–*i*.*e*., residence since birth or childhood. For that reason, after characterizing the genetic structure and ancestry of the studied population, we aim to conduct transcriptome analyses of blood cells in Peruvian individuals, primarily from rural areas, from indigenous populations. These individuals are exposed to varying living altitudes, comparing individuals from high versus low altitudes. We collected peripheral blood mononuclear cells (PBMC) from 62 volunteers, who were exposed to bacterial lipopolysaccharide (LPS), Pam3CSK4 (a synthetic triacylated lipopeptide responsible for bacterial components), and R848 (an imidazoquinoline compound related to viral nucleic acids) (14). Subsequently, we discerned variations in gene expression profiles among populations residing at low and high altitudes (hypoxia). Our analysis primarily concentrated on genes that exhibited differential expression regardless of stimuli, with a special focus on those significantly involved in immune responses.

## MATERIAL AND METHODS

### Population and samples

We studied the population from Huanuco, a department in Central Peru, focusing on a region in the eastern slope of the Andes that includes the Andean Mountains and the Amazonian Tropical Forest (in particular the so-called Amazon Yunga or Higher Amazonia). In addition to biological samples, we collected demographic data such as sex, age, body mass index (BMI), and altitude of residence. In the first part of the study, 35 individuals from the studied population were analyzed for genomic diversity. In the second part of the study, we studied 62 individuals for transcriptome analyses, with an overlap of 19 individuals between the two parts of the study. The study was approved by the Inmensa Ethics Committee of Peru (number 0011-2021). Written informed consent was obtained from all participants involved in this study.

### Genetic structure and ancestry of the studied Huanuco population

#### Genotyping and Data Quality Control (QC)

Thirty-five individuals were genotyped with the Illumina GSA MG v3 array by Macrogen (Seul, Korea). After Illumina GenomeStudio processing, we obtained 737,534 SNVs. After further QC with MosaiQC software (https://github.com/ldgh/Smart-cleaning-public), a tool that automates the Laboratory of Human Genetic Diversity (LDGH) quality control pipeline using PLINK, 703,515 SNVs remained.

Individual ancestry analyses. For the genetic clustering analysis, we ran ADMIXTURE (15), which estimates the fraction of each K parental population that contributes to an individual by fitting the Hardy-Weinberg equilibrium in each of the K populations/clusters (7), using 189,837 unlinked SNVs (r^2^<0.4). To cover the admixture history and the structure inside Peruvian Native populations, we first ran 14 replicates in the unsupervised mode for K=3 to infer African, European and Native American ancestry using the entire dataset in Supplementary Table S1. We presented the replicate with the higher log-likelihood value.

To avoid the bias inserted by related individuals in ADMIXTURE inferences, we used K=3 to estimate kinship coefficients between Huánuco individuals (Φij) with the software REAP (Relatedness Estimation in Admixed Populations), which is more appropriate for admixed individuals (16). The estimates of Φij by REAP are conditioned on the estimated individual ancestry proportions (estimated with ADMIXTURE) for K=3 parental populations. To identify related individuals to be pruned for further ancestry analyses (K>3), we used NAToRA, a relatedness-pruning method to minimize the loss of dataset in genetic and omics analyses (17), using the relatedness cutoff of Φij > 0.06.

Then, we pruned our dataset for the few related individuals and ran again 14 replicates of ADMIXTURE for K from 4 to 12, considering only unrelated individuals and 162,957 unlinked SNVs, presenting the results of the most likely run for each K, and assessing CV errors for different K. Because the ADMIXTURE results with K=3 informed us that our dataset had negligible African ancestry, we removed African individuals from the ADMIXTURE analyses for K>3.

### Transcriptome analyses in the Huanuco population

#### Preparation of biological samples

In 62 individuals, PBMCs were isolated from whole blood by Ficoll-Paque centrifugation within 4 hours of collection and cryopreserved. PBMCs were cultured in RPMI-1640 (Fisher) supplemented with 10% heat-inactivated FBS (FBS premium, US origin, Wisent) and 1% L-glutamine (Fisher). All samples were stimulated by pathogen molecules (virus and bacteria) in independent processes: for each of the tested individuals, and one process without stimulation.

Bacterial and Virus Stimulation Assays PBMCs (2 million per condition) were stimulated for 4 hours at 37° C with 5% CO2 with bacterial lipopolysaccharide (LPS), Pam3CSK4 (Invitrogen – tlrl-pms) a synthetic triacylated lipopeptide responsible for bacterial components, and R848 (Invitrogen – tlrl-r848-5) an imidazoquinoline compound, related to viral nucleic acids, according to the manufacturer’s instructions.

RNA Extraction Total RNA was extracted with the PureLink RNA Mini Kit from Invitrogen Inc., including the enzymatic digestion of genomic DNA. Extractions were performed in batches of 10 samples, and RNA quality and quantity were assessed with a Nanodrop spectrometer and the Agilent Bioanalyzer RNA 6000 nano kit. We generated a final set of 79 samples from the 62 donors for high-throughput RNA-sequencing, including 17, 16, 24, and 22 samples for the non-simulated, LPS, Pam3CSK4, and R848 conditions, respectively.

#### RNA Library Construction, Quality Control, and Sequencing

The process of library construction involved the purification of messenger RNA from total RNA using magnetic beads attached to poly-T oligos (TruSeq RNA Sample Prep Kit v2, Illumina Inc.). Subsequently, the mRNA was fragmented, and the first strand of cDNA was synthesized using random hexamer primers. This was followed by the synthesis of the second strand cDNA. The library was considered complete after undergoing end repair, A-tailing, adapter ligation, size selection, amplification, and purification. Quantification and size distribution analysis of the library was performed using Qubit and real-time PCR, as well as a Bioanalyzer (Agilent, CA, USA). Finally, the pooled libraries were sequenced using Illumina platforms by the NOVAGENE facility (https://www.novogene.com/). The total data output sequencing product was 514.1 GB. The Data Quality report and summary are available in the Supporting Information and Data.

#### Transcriptome data pre-processing

Total RNA sequencing was performed by a pair-end strategy for all samples using Novaseq PE150 (Illumina Inc). Raw data is stored in .fastq format (.fq.gz compressed) for analysis. The NOVOGENE facility provides raw RNA sequencing data by demultiplexing (separating multiple samples pooled into a single library), removing adapters, and low-quality reads regarding bases that cannot be determined (N > 10%) and those with QScore <= 5. For our quality control analysis, each sample in .fastq was resubmitted for the quality control pipeline using fastQC and multiQC to generate sequencing quality reports to verify quality per bases, GC content, the proportion of undefined nucleotide bases, and the presence of adapter sequences. The Phred score Q30 was the base threshold for filtering high-quality samples. Read alignment was performed with STAR with the Release 41 (GRCh38.p13) genome reference. The software STAR was set to consider that a read may overlap on the same or opposite strand and to follow the Encyclopedia of DNA Elements (ENCODE) best practices. The STAR outputs alignments in a sorted BAM format. After genome alignment, read quantification was performed with HTSeq with the GRCh38 human genome annotation file (.gff). The read-counting process considered the large set of transcripts, in total 61801 coding and non-coding elements cataloged in ENCODE. To avoid non-expressed transcripts, we used the *filterByExpr()* function to filter for genes that meet a minimum expression threshold. In this analysis, only genes with at least 10 reads across all samples are retained.

#### RNA-seq variant calling and global ancestry analysis

Some individuals were sequenced for RNA-seq multiple times under different stimuli; thus, we merged the corresponding FASTQ files for each individual. We then used GATK’s Best Practices pipeline (https://github.com/gatk-workflows/gatk4-rnaseq-germline-snps-indels), incorporating steps for joint variant calling. Variants were filtered using the recommended parameters: a minimum depth of 5 and up to 10% missing data, resulting in 85,557 variants. For ancestry inference, we selected 11,861 unlinked SNPs (r^2^ < 0.4) and performed ADMIXTURE analysis (Alexander, Novembre, and Lange, 2009) in unsupervised mode, testing K values from 3 to 12 with three replicates each. African individuals were excluded from panels for subsequent ADMIXTURE analyses, due to African ancestry showing negligible contributions. Using the resulting set of variants and quality control filters, we conducted principal component analysis (PCA) using EIGENSOFT (18,19) to compute 12 principal components.

#### Differential gene expression analysis

The 17 control samples (i.e. not stimulated) and 62 treated PBMC samples LPS (n = 16), Pam3CSK4 (n = 24), R848 (n = 22)) were also organized in two groups: low altitude (less than 1000 m.a.s.l.) and high altitude (>3000 m.a.s.l.) to perform differential expression analysis (DEA) with the edgeR package (20)

Gene expression variance was estimated by variance partition using a linear mixed model (21). Furthermore, the dispersion of gene expression was assessed to fit a generalized linear model including covariate correction for individual (categorical variable), sex (categorical variable), BMI (numerical variable), age (numerical variable), stimulus by pathogens (one virus and two bacteria, categorical variable), and Native ancestry proportion. Those genes with a false-discovery rate (FDR < 0,05) and fold-change (|FC| >= 2) were considered differentially expressed genes (DEG). The distribution of DEG genes (down- and up-regulated) regarding statistical significance and genes from the immune system, before and after stimulation, was visualized using volcano plots and heatmaps, created with the EnhancedVolcano (https://github.com/kevinblighe/EnhancedVolcano) and pheatmap packages in R (https://cran.r-project.org/package=pheatmap). Hierarchical clustering of genes and samples was performed and visualized on heat maps using Pearson correlation and the complete linkage method (19, 20).

#### Functional Enrichment Analysis

We conducted a thorough functional enrichment analysis of gene sets, encompassing both up-regulated and down-regulated genes, employing Gene Ontology (GO) and utilizing the ClusterProfile (22) package in the R language. To enhance statistical rigor, multiple tests were corrected using FDR. The outcomes, comprising the top biological processes, were visually represented through bar plots and networks depicting the interrelation of genes and processes.

#### Immune system genes dataset

DEA results were merged with biological pathway data, focusing exclusively on genes related to the Immune System. In the Reactome database, the adaptive *immune system* catalogs the interaction of 868 genes, the *innate immune system* shows 724 transcripts, and 1160 genes comprise the *cytokine signaling pathway*. The Reactome data is available at reacomte.org, and we used version 81 released on June 16, 2024.

## RESULTS

### Genomic diversity and ancestry of the Huanuco population

ADMIXTURE analyses (Figure 1, K=3 and 4) of 35 individuals reveal that Huanuco individuals are predominantly Native American (87.3% on average) with a low European ancestry (average: 11.82%) and negligible African ancestry (<1%) (Supplementary Table S2). Given the negligible African ancestry, which is consistent with the history of the Huanuco region, we present ADMIXTURE results excluding African individuals (results in Supplementary Table S3).

**Figure 1.**
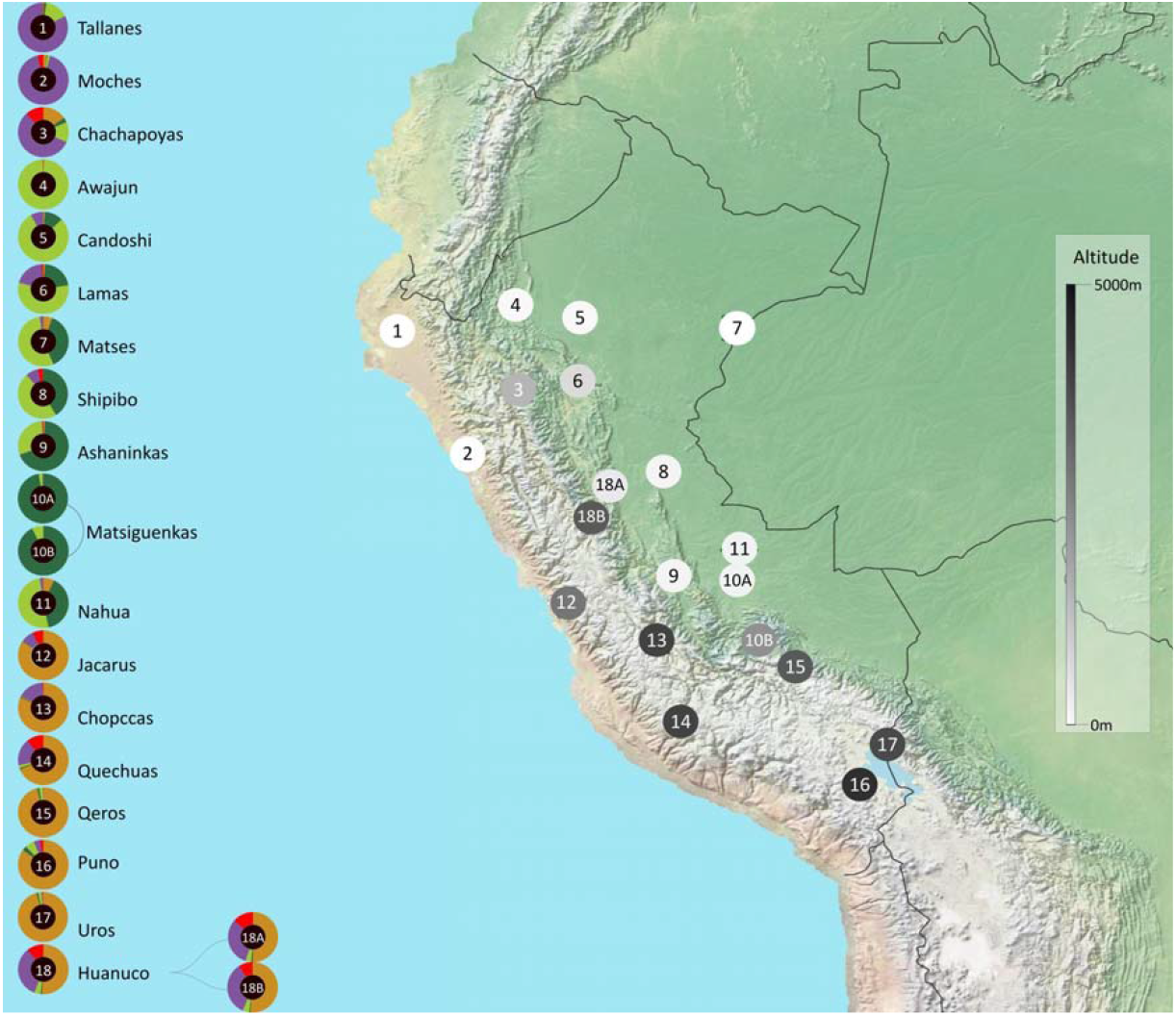
Genetic and geographic landscapes for Peruvian populations. (A): Genetic structure of seventeen native populations from the coast (n. 1 and 2), Amazon (n. 3-11), Andes (n. 12-17), and Huanuco samples (18A and 18B), inferred by ADMIXTURE (K=6). Donut plots show the average percentages of individuals for six ancestry clusters in each population. Four clusters were associated with distinct Native American groups: one Andean (brown), one North Amazonian (light green), one South Amazonian (dark green), and one coast (purple). Two clusters were associated with non-Native continental ancestries, both European (red), caused by admixture with these populations in the last 500 years. (B): Information for geographic location and altitude of samples. The color of the population markers goes from white (at sea level) to black (5000 meters above sea level). (https://www.naturalearthdata.com/)

Analyses assuming K=5 ancestry clusters (K=4 and 5) reveal that Native American ancestry of Huánuco individuals may be dissected in a predominant Andean-associated cluster (84.5% [SD:5.6%] of the total ancestry) and a minor Amazonian-associated cluster (3.7% [SD: 1.1%] of the total ancestry). Interestingly, the Andean-associated ancestry cluster is higher in highlanders (average 86%) than in lowlanders (average 82%, <600 m vs. >3000m, Kruskal-Wallis H=4.057, p<0.05, N=35).

A higher-resolution clustering (K = 6 and 7) reveals that Huanuco individuals are similar to Tallanes and Moche (indigenous groups from the northern Peruvian Coast) and Chacapoyas (an indigenous group from the north of the Andes/Amazonian geographic interface). Altogether, Huanuco individuals are similar to Tallanes, Moche, and Chachapoyas from Northern Peru, and all of them present different levels of admixture between Andean and Amazonian ancestries.

### Demographic data overview of the Huanuco transcriptome study

Forty-three (69.4% or total) participants were highland sojourners (3226 – 3539 m.a.s.l.), whereas 19 (30.6%) inhabited lowlands (596 – 634 m.a.s.l.). The samples were obtained at the residences of participants and comprised 11 males (11.7%) and 51 females (82.3%) with an age mean of 48 years (SD±15). The samples showed a BMI median of 26.4 Kg/m^2^. It is worth noting that almost all participants (high and lowlanders) had ancestors (e.g., parents, grandparents) from high Andean areas of the Huánuco department (Table 1).

**Table 1.** Demographic data of Peruvian individuals from rural areas, categorized by different living altitudes (high vs low).

| Variables | Total |  | Living altitudes |  |  |  | p-Value |
| --- | --- | --- | --- | --- | --- | --- | --- |
|  | N | % | Low |  | High |  |  |
|  |  |  | N | % | N | % |  |
| **Sex** |  |  |  |  |  |  |  |
| Female | 51 | 82.26 | 15 | 78.95 | 36 | 83.72 |  |
| Male | 11 | 17.74 | 4 | 21.05 | 7 | 16.28 | 0.650 |
| **Age (years)** | - | - | 42 | (34 - 58) † | 48 | (37 - 64) † | 0.392 |
| **Cholesterol (mg/dl)** | - | - | 169 | (136 – 185)† | 166 | (145 – 204)† | 0.262 |
| **Hemoglobin (g/dl)** | - | - | 14.4 | (13.4 - 15.1)† | 14.1 | (13.4 - 14.4)† | 0.267 |
| **Glucose (mg/dl)** | - | - | 87 | (78 - 96)† | 83 | (76 – 89)† | 0.184 |
| **BMI (kg/m<sup>2</sup>)** | - | - | 28.3 | (24.3 - 30.2)† | 25.1 | (22.3 - 28.5)† | 0.031 |

\*Median. †(Q1 - Q3)
BMI (Body mass index)
Statistically significant ( $p < 0.05$ , Chi-squared).

### Differentially expressed immune genes in lowlanders versus highlanders

After applying the count filtering criterion, the number of transcripts is reduced from 61,806 to 14,832, effectively avoiding transcripts inflated with low expression (counts < 10) per sample. After quality control including contamination check, the distribution of samples stratified for stimulation and altitude is detailed in Supplementary Table S4.

Gene expression variance is primarily driven by individual-specific differences (21.80%), followed closely by stimulus group effects (16.99%). Moderate contributions come from altitude (5.45%) and age (2.49%). Smaller but significant influences include population-specific variation (1.71%), sex differences (1.54%), and body mass index (1.47%). Together, these factors highlight the complex interplay of genetic, environmental, and demographic factors shaping gene expression variability (Supplementary Figure S2).

Comparative analysis between individuals from low and high altitudes revealed 31 differentially expressed genes (DEGs), accounting for covariates such as individual differences, sex, BMI, age, Native American ancestry proportions, and pathogen exposure (both viral and bacterial). Among these DEGs, 22 were up-regulated and nine were down-regulated in high-altitude individuals compared to those at low altitude. Notably, three immune-related genes (*FN1, CD36, FOS*) were significantly down-regulated in high-altitude individuals (Figure 2A, Table 2). All DEGs are listed in Supplementary Table S5.

**Table 2.**
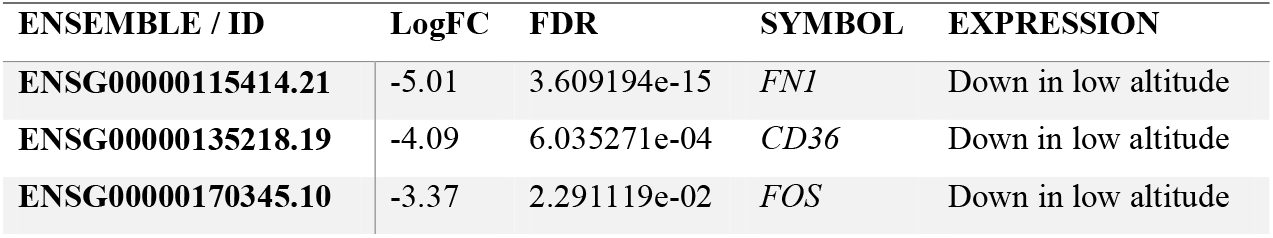
Differential gene expression analysis results.

**Figure 2.**
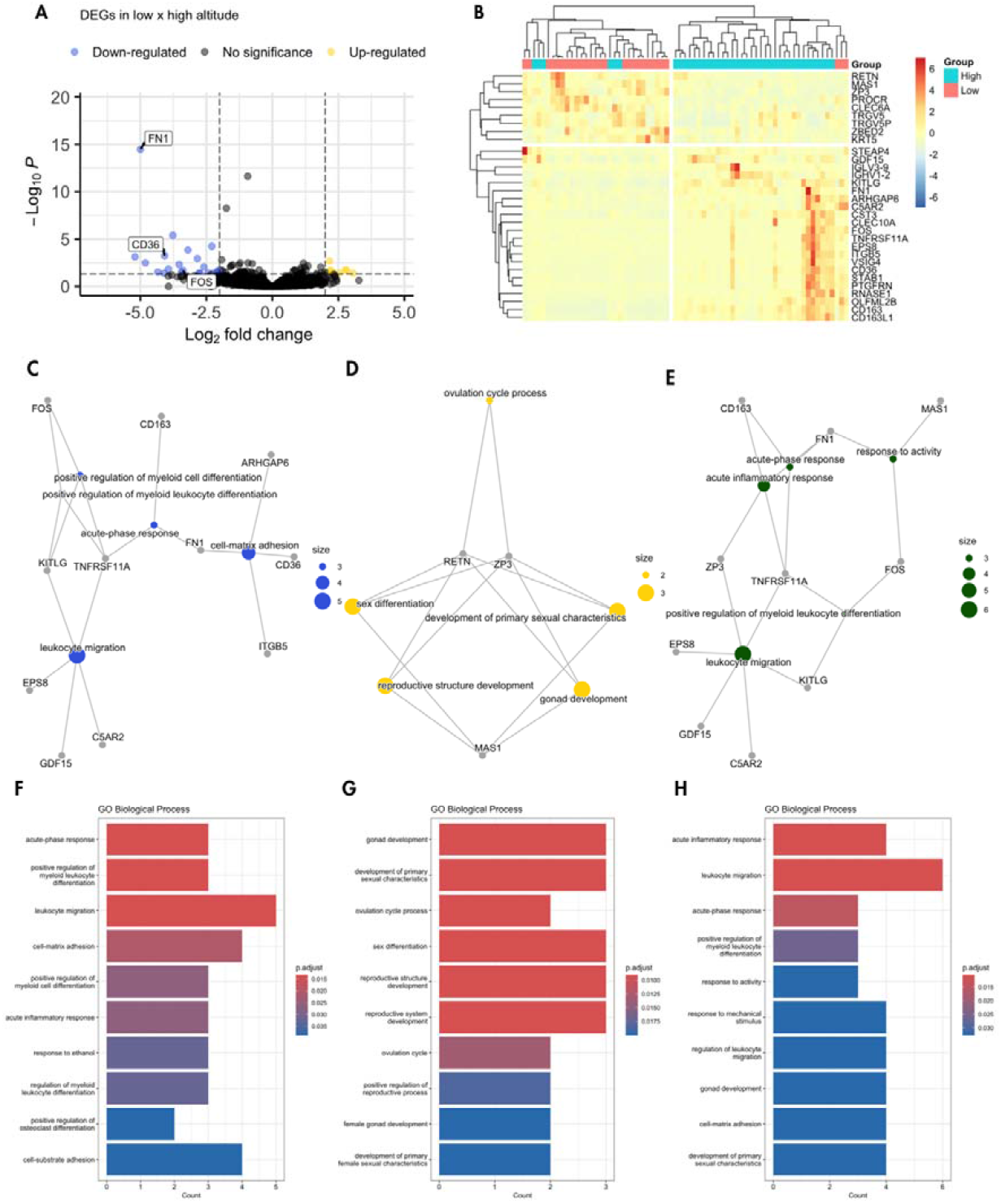
Differentially Expressed Genes Analysis: A) The volcano plot illustrates the most significantly expressed immune system genes between low and high altitude residents. Genes are plotted based on their expression levels and statistical significance (P ≤ 0.05, FC ≥ 2), highlighting three down-regulated *FN1, CD36, and FOS*. B) Hierarchical clustering of differentially expressed genes (DEGs) and samples is shown, with gene expression values normalized by Z-score. Functional enrichment analysis of Gene Ontology was performed using ClusterProfile’s enrichGO function, focusing on biological processes for the entire set of DEGs (C, F), down-regulated genes (D, G), and up-regulated genes (E, H). The x-axis represents the number of genes associated with each biological process, and the false discovery rate (adjusted p-values) is indicated by a red-blue gradient, showing statistical significance across different methods. P: False Discovery Rate (adjusted p-values); FC: Fold change.

We performed hierarchical clustering on DEGs identified between lowlanders versus highlanders samples (Figure 2B). The resulting dendrogram revealed two distinct clusters of genes. Furthermore, the sample dendrogram showed a separation between lowlander and highlander groups, indicating distinct gene expression profiles. The heatmap visualization, with dendrograms overlaid, provided a detailed view of the expression patterns, highlighting key clusters of DEGs. Despite accounting for covariates like sex, BMI, age, Native American ancestry proportions, and pathogen exposure, there might be additional unmeasured factors influencing gene expression, causing some samples to cluster differently.

Functional enrichment by gene ontology analysis sheds light on the molecular landscape underlying cellular responses, particularly in the context of immune and inflammatory processes (see Figure 2C-H). The complete functional enrichment analysis results are listed in Supplementary Table S6-S8.

The downregulated genes show a strong enrichment for immune and inflammatory processes, including acute-phase and acute inflammatory responses, leukocyte migration, and myeloid lineage differentiation. Several terms also relate to cell adhesion, suggesting a potential decrease in cell–matrix or cell–substrate interactions. Together, these patterns indicate a suppression of inflammatory signaling and myeloid cell activity, which could reflect reduced immune cell infiltration and altered immune responsiveness.

In contrast, the upregulated genes are enriched for reproductive and developmental processes, particularly those associated with gonad development, ovulation cycle regulation, and sexual differentiation. Many terms are specific to female reproductive biology, indicating activation of hormone-driven pathways such as estrogen or progesterone signaling. This transcriptional shift from immune-related programs toward reproductive development could be driven by physiological states, hormonal stimuli, or experimental conditions that favor reproductive tissue function over immune activity.

Cell-type-specific Enrichment Analysis (CSEA) for differentially expressed genes was conducted using WebCSEA, a web-based tool designed for Cell-type Specific Enrichment Analysis of Genes. By leveraging WebCSEA, we found cell-type-specific gene expression profiles for macrophages and dendritic cells (Supplementary Figure S3).

## DISCUSSION

Our study is the first to investigate the gene expression of indigenous populations from Peru (Figure 1) concerning their immune response, considering the altitude of their residence, which generally exposes them to low oxygen concentration, cold, dryness, ultraviolet radiation, and a differentiated immune response. In this sense, Quach et al. recently reported that the effect of immunity may be related to ancestry, which an individual or population belongs to; specifically, Africans tend to have a stronger immune response than Europeans (14).

This is the first study to evaluate differential immune response in a high-altitude population with high native ancestry, where we have identified DEGs. The downregulation of *FN1, CD36, and FOS*, involved in processes related to extracellular matrix, cell adhesion, and immune response, could improve our understanding of the modulation of the innate and adaptive immune systems. Our results are congruent with what has been observed in previous research conducted on Andean populations, which have identified genes involved in the immune system, such as *DUOX2, CD40, TNIP1*, and *CHIA* (7,11).

A recent study published by Li *et al*. presents a meta-analysis of immune genes that investigated how they varied in expression when exposed to altitude conditions, concerning studies of cells exposed to normoxia or hypoxia. In the bioinformatics analysis, the cells came from European descent subjects exposed to altitude but not native residents of a high-altitude area (23). Li *et al*. demonstrated that the genes *ERH, VBP1, PSMA4, TOMM5*, and *POLR2K* were downregulated in association with hypoxia; however, our results show that these genes are not statistically differentiated. Additionally, these authors show that altitude promotes the increase of some immune cells (T cells CD4 memory resting, T cells gamma delta, Dendritic cells activated, Mast cells resting) at the expense of others relevant to the innate or adaptive immune system (*e*.*g*., T cells CD8, NK cells resting, Neutrophils).

In contrast, Manella *et al*. discovered that genes associated with immune response can exhibit differential expression patterns based on both the duration of acute exposure to high altitude and the specific altitude level (e.g., 3,800 meters vs. 5,100 meters). Additionally, the timing of the measurements during the day also plays a role in these gene expression variations (24). The investigation revealed that time-associated gene alterations were linked to innate immunity, specifically about the activation of neutrophils. Additionally, distinct transcriptional patterns were identified concerning the factors *RELA* and *IRF8*, recognized as mediators of inflammation and immune responses. Furthermore, the study conducted by Manella et al. unveiled an elevation in M2 macrophages and monocytes, influenced by both altitude and time. Conversely, a decrease contingent on altitude was observed in NK cells and regulatory T cells. Moreover, the researchers demonstrated the temporal variation in the expression of *NR1D1*, with its overall expression being notably reduced in response to altitude (25).

The Manella team’s investigation is particularly intriguing due to its unique focus on individuals of European descent in one of the world’s most densely inhabited cities, located in Peru. However, it focused on European descent men subjected to a relatively short-term exposure, spanning several weeks, which cannot be directly compared to the chronic exposure experienced by native inhabitants at such altitudes. Moreover, our study population in Huánuco regularly experiences altitude fluctuations due to the necessity for individuals residing in the high Andean regions to descend to the city of Huánuco for various activities, such as shopping, making payments, receiving government assistance, and engaging in recreational pursuits, among others. As a result, a prospective study conducted within the Huánuco population, using a sample resembling ours, could investigate how genes are regulated, either up-regulated or down-regulated, with the duration of absence from high-altitude environments followed by return or conversely, the effects of prolonged high-altitude exposure and subsequent return to lower altitudes.

Thus, we can assert that our study has demonstrated a distinct genetic expression signature in high-altitude residents with a native Andean genetic background, as compared to their counterparts living at lower altitudes. Furthermore, we have identified several genes exhibiting both up-regulation and down-regulation, with a distribution observed within genes related to the innate and adaptive immune systems.

It’s worth emphasizing that most of our study participants were female, and we did not implement age-matching procedures during the sample selection process. This absence of age matching could potentially introduce selection bias, as gene expression can vary based on both the participant’s sex and age. To mitigate potential sex-related bias, we incorporated sex as a covariate variable in our differential expression analysis. However, we acknowledge that our study was limited in its ability to consider all environmental conditions and travel habits, factors that may have exerted an influence on our research findings.

A significant limitation of this study on the genomics and transcriptomics of lowlander and highlander populations is the small sample size. With few samples, the ability to generalize the results to the entire population is limited. Additionally, the small sample size increases susceptibility to statistical biases and reduces the statistical power to detect subtle differences or significant biological effects. However, our study stands as the pioneering effort demonstrating how immune response gene expression varies in individuals chronically exposed to high altitude (natives). Further investigations could validate and expand upon our findings through genomics, including transcription factors, transcriptomic, proteomic, or even epigenetic analyses.

## CONCLUSIONS

The deregulated genes *FN1, CD36*, and *FOS* in native inhabitants with a prominent Andean genetic background highlight the active involvement of both the innate and adaptive immune systems. Remarkably, our study suggests that the immune response, even if induced by the same stimulus, varies with altitude. A comprehensive multi-omic approach is indispensable to fully understand the impact of altitude on immune system gene expression and genes related to high-altitude adaptation.

## ACKNOWLEDGEMENTS

We thank Lluis Quintana-Murci and David Comas for their constructive comments and perspectives. We thank all those who facilitated the recruitment of participants, including the Direccion Regional de Salud from Huanuco; and all participants in this study.

## SUPPORTING INFORMATION

**Figure S1. Huanuco population structure based on the subset of samples with intersecting genomic and transcriptomic data.**

**Figure S2. Variance Partition Analysis.**

**Figure S3. Cell-type-specific Enrichment Analysis (CSEA) for differentially expressed genes was obtained using WebCSEA (Web-based Cell-type Specific Enrichment Analysis of Genes).**

**Table S1 - Population data**

**Table S2 - Population ancestry**

**Table S3 - Huanuco ancestry data**

**Table S4 - Distribution of samples per group and altitude**

**Table S5 - All DEGs**

**Table S6 - Functional enrichment analysis for down-regulated genes**

**Table S7 - Functional enrichment analysis for up-regulated genes**

**Table S8 - Functional enrichment for all deregulated genes**

**File: Analysis scripts File: Quality Control**

---

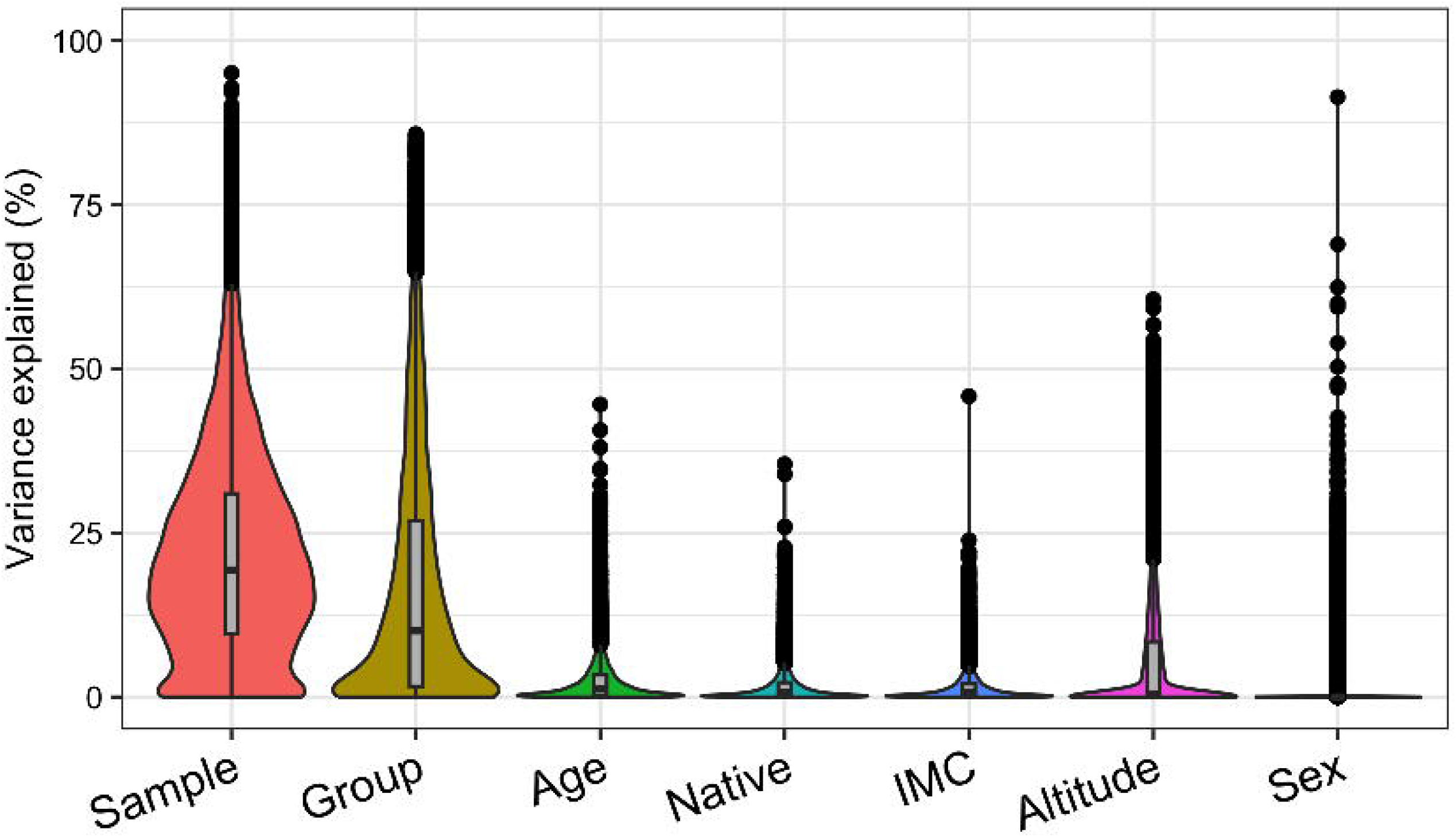

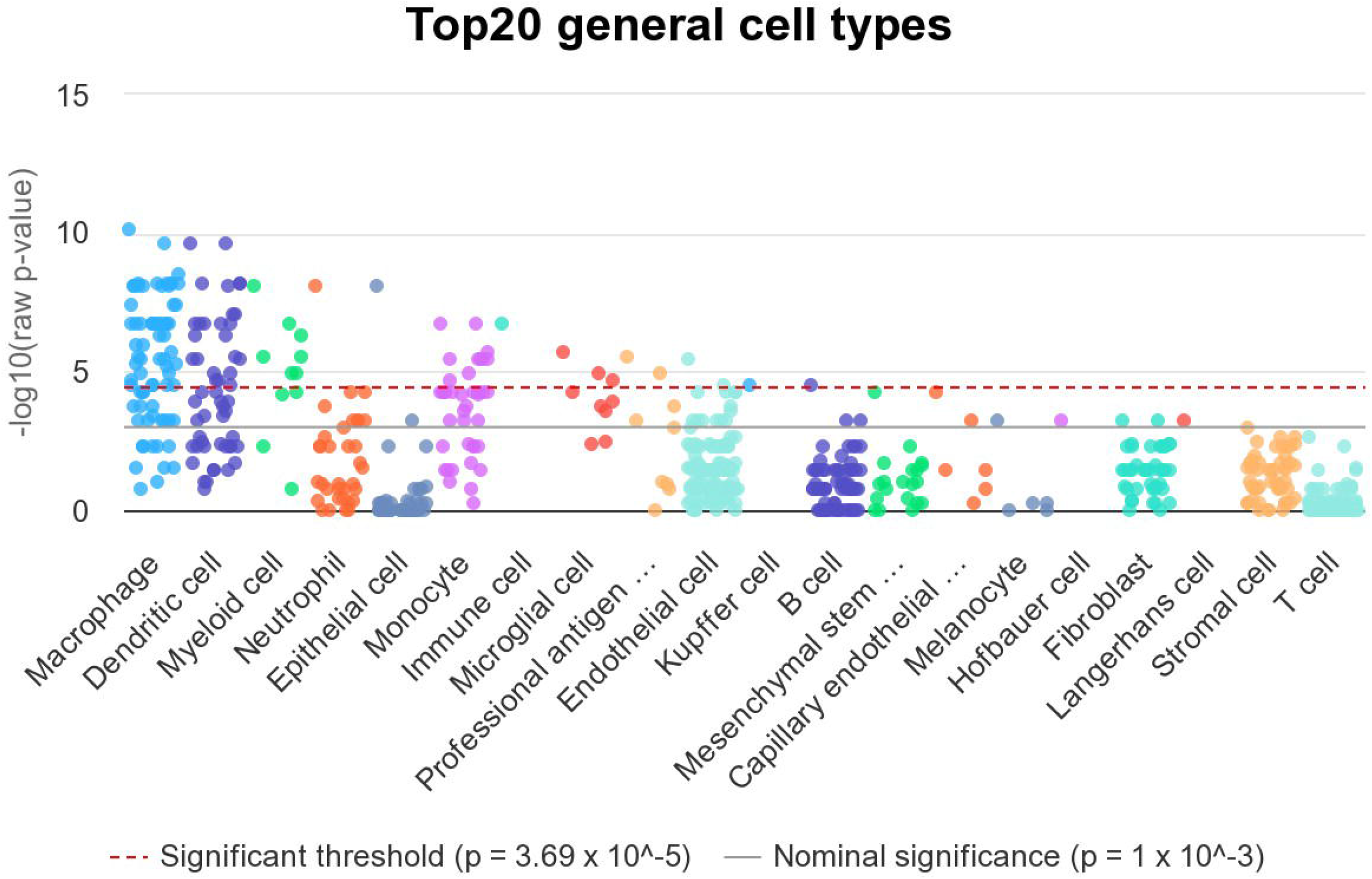

